# Functional plasticity of AIF revealed by dimerization and CHCHD4 interaction states

**DOI:** 10.64898/2026.08.31.748248

**Authors:** Olga Soriano, Soraya Hernández-Hatibi, Rebeca Gracia-Domingo, Silvia Romero-Tamayo, Miguel Ferrer, Adrián Velázquez-Campoy, Joaquín Marco-Brualla, Patricio Fernández-Silva, Santos A. Susin, Milagros Medina, Raquel Moreno-Loshuertos, Patricia Ferreira

## Abstract

Apoptosis-inducing factor is a mitochondrial flavoprotein that links redox metabolism to mitochondrial homeostasis through its interaction with the disulfide relay protein CHCHD4. Although NADH-dependent AIF dimerization has been proposed as the activated state mediating CHCHD4 engagement, whether it is strictly required for productive AIF–CHCHD4 function remains unclear.

Here, combining cellular, biochemical and biophysical approaches, we show that disruption of the AIF dimer interface compromises oxidative phosphorylation, respiratory-chain organization and CHCHD4-dependent mitochondrial homeostasis, yet preserves partial AIF function. Our data reveal that the AIF–CHCHD4 system operates as a conformational dynamic redox module in which distinct AIF oligomeric and redox states sustain CHCHD4 activity with different efficiencies. Mechanistically, dimerization is coupled to NADH-dependent conformational changes that regulate coenzyme binding, charge-transfer complex stabilization and catalytic efficiency. In turn, CHCHD4 binding remodels AIF conformational and redox properties, partially compensating for defects in dimer stabilization or redox coupling. Consistently, a peptide derived from the CHCHD4 N-terminus partially restores redox function in a pathogenic AIF variant defective in dimer stabilization, supporting partner-assisted allosteric regulation as a potential therapeutic strategy.

## Introduction

The human apoptosis-inducing factor (hAIF) is a mitochondrial flavoenzyme that contributes both to cell death, through the caspase-independent pathway known as Parthanatos, and to cell survival (Joza *et al*, 2001; Susin *et al*, 1999). In healthy cells, hAIF is located in the mitochondrial intermembrane space (IMS), where it is anchored to the inner membrane and contributes to mitochondrial architecture and redox homeostasis (Wischhof *et al*, 2022; Xie *et al*, 2005). Upon apoptotic stimuli, AIF translocates to the nucleus and promotes poly(ADP-ribose)polymerase (PARP)-1-dependent chromatinolysis through the formation of a DNA-degradosome complex (Alano *et al*, 2010; Novo *et al*, 2023; Wang *et al*, 2016). Beyond its apoptotic role, increasing evidence supports a function for AIF as a redox-responsive regulator of mitochondrial protein homeostasis.

Structurally, hAIF contains FAD– and NADH-binding domains responsible for its oxidoreductase activity, and a C-terminal domain involved in protein-protein interaction and apoptotic signaling (Fig. 1A). Reduction of the FAD cofactor by NADH induces conformational rearrangements at the active site associated with the formation of a stable FADH⁻/NAD⁺ charge-transfer complex (CTC), remodeling of regulatory regions including the C-loop (residues 509-560) release, and exposure of a hydrophobic surface that promotes AIF dimerization (Brosey *et al*, 2016; Ferreira *et al*, 2014; Romero-Tamayo *et al*, 2021; Sevrioukova, 2009) (Fig. 1B). The dimer is stabilized by a network of interactions involving residues E413, R422 and R430 (Ferreira *et al*., 2014). A central element in this process is residue H454, which participates in the redox-linked conformational network coupling CTC stabilization to NADH-dependent structural rearrangements and dimer formation (Brosey *et al*., 2016; Villanueva *et al*, 2015). Recent structural studies further indicate that these NADH-dependent conformational transitions generate a remodeled C-terminal interaction platform that mediates binding to partner proteins such as CHCHD4 (coiled-coil-helix-coiled-coil-helix-domain containing 4; Mia40 in yeast), linking redox sensing to mitochondrial protein homeostasis (Brosey *et al*, 2025; Rothemann *et al*, 2025). The structural basis of NADH-dependent allosteric dimerization of AIF and the role key residues are described in detail in Supplementary information.

**Figure 1.**
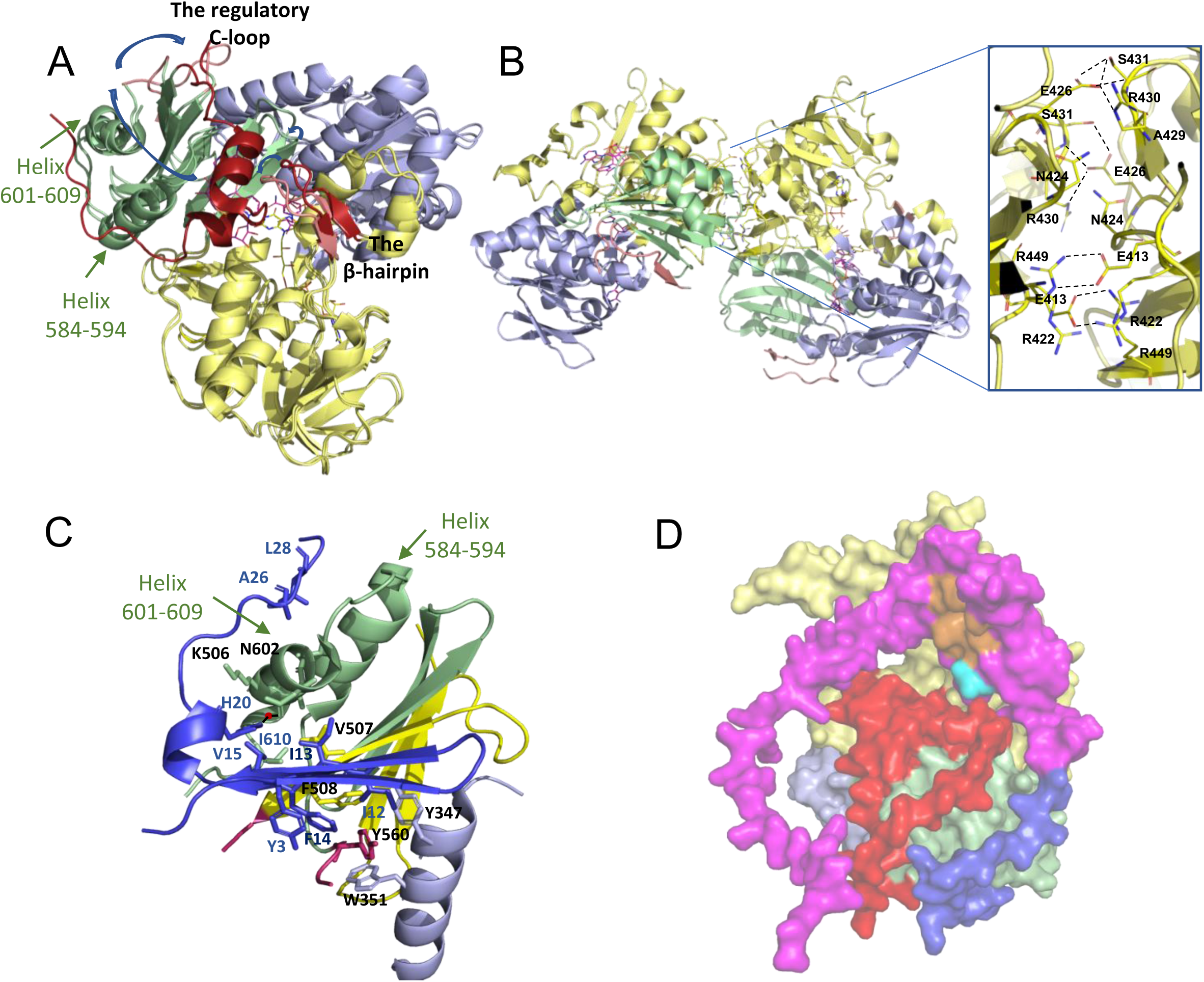
– Structural insights into human AIF redox dependent conformational changes, dimerization, and interaction with CHCHD4. **A** Overlay of oxidized and one of the monomers reduced WT AIF (chains A from PDB entries 4BV6 and 4BUR respectively) in cartoon representation. **B** Cartoon representation of the WT AIF_Δ101rd_:2NAD^+^/H CTC homodimer. A zoomed view of the dimerization interface (right) highlights key residues as CPK-colored sticks, with interchain polar interactions indicated by dashed lines. **C** Detail of the complex between monomeric hAIF W196A_Δ127_ and the N-terminal peptide of CHCHD4 (PDB 8VGY) in cartoon representation. The N-terminal CHCHD4 is coloured in blue. Key binding residues in AIF and CHCHD4 are shown as sticks, labelled in black and blue, respectively. **C** Surface representation of the AlphaFold model of the AIF:CHCHD4 complex, as extracted from(Pei *et al*., 2022). The C-terminal region of CHCHD4 is shown in pink, and its N-terminal domain is in blue. Hydrophobic residues involved in CHCHD4 substrate recognition are highlighted in orange, and the CPC motif in cyan. In all panels, FAD-binding (residues 130–262 and 401–480), NADH-binding (residues 263-400), and C-terminal (residues 481-613) domains of AIF are coloured in yellow, light blue and lime green, respectively. The regulatory C-loop (residues 505-560) and the β-hairpin (residues 190-202) are respectively highlighted in red and salmon in oxidized and reduced structures. FAD and coenzymes (NAD(H)_A_ and NAD(H)_B_) are displayed as CPK-coloured sticks, with their C atoms in yellow and pink, respectively.

In the IMS, AIF exists in a monomer-dimer equilibrium and contributes to mitochondrial redox metabolism through its interaction with CHCHD4, a central component of the disulfide relay system (DRS) responsible for oxidative folding of assembly factors and respiratory-chain subunits (Hangen *et al*, 2015; Salscheider *et al*, 2022). CHCHD4 catalyzes disulfide bond formation in cysteine-rich proteins through its redox active cysteine-proline-cysteine (CPC) motif and is reoxidized by the sulfhydryl oxidase ALR (augmenter of liver regeneration) to sustain the DRS cycle (Reinhardt *et al*, 2020). This pathway is essential for respiratory-chain biogenesis and mitochondrial organization, placing CHCHD4 at the center of IMS proteostasis.

AIF has been proposed to regulate CHCHD4 through a NADH-dependent complex (Hangen *et al*., 2015). Recent structural studies show that the N-terminal region of CHCHD4 contains an AIF-interaction motif that mimics unfolded mitochondrial import substrates, allowing CHCHD4 to engage a hydrophobic cavity within the AIF C-terminal domain that becomes exposed upon the NADH-induced conformational remodeling (Fig. 1C-D) (Brosey *et al*., 2025; Fagnani *et al*, 2024; Pei *et al*, 2022). Cryo-EM structures further demonstrate that CHCHD4 binding stabilizes the reduced AIF CTC dimer while propagating conformational rearrangements back to the active site (Rothemann *et al*., 2025). Together, these observations support a model in which AIF dynamically coordinates CHCHD4 activity and mitochondrial redox homeostasis through conformationally coupled protein–protein interactions.

Defects in either AIF or CHCHD4 impair OXPHOS and compromise mitochondrial function, highlighting the physiological relevance of this pathway (Hangen *et al*., 2015; Meyer *et al*, 2015). Consistently, CHCHD4 overexpression partially compensates for AIF deficiency in cells. Moreover, more than 20 mutations in the hAIF gene (*AIFM1*) have been associated to rare neurodegenerative disorders with highly heterogeneous phenotypes (Wischhof *et al*., 2022). Several disease-associated variants alter CTC stability and OXPHOS activity, suggesting that perturbation of AIF conformational dynamics and CHCHD4 interaction contributes to mitochondrial dysfunction (Ferrer *et al*, 2026; Qiu *et al*, 2023; Sevrioukova, 2016; Sorrentino *et al*, 2017). However, despite recent structural advances, it remains unclear whether NADH-induced AIF dimerization is strictly required for CHCHD4 regulation and mitochondrial homeostasis, or instead represents a highly efficient but non-obligatory functional state.

Here, we combined biochemical, biophysical and cellular approaches to define the contribution of AIF dimerization to CHCHD4 interaction and mitochondrial redox regulation. We characterized two separation-of-function AIF variants: H454A, which adopts a dimer-permissive conformation while retaining an oxidized-like C-loop organization in both redox states (Brosey *et al*., 2016); and E413A/R422A/R430A, which fails to stabilize the NADH-induced dimer (Ferreira *et al*., 2014). We further generated a cellular model harboring the E413A/R422A/R430A substitutions to assess the impact of impaired dimerization on OXPHOS and CHCHD4 interactions. Finally, we explored the ability of a CHCHD4-derived peptide to modulate the pathogenic R422Q variant displaying defective dimer stabilization (Qiu *et al*., 2023). Our findings reveal a previously unrecognized functional plasticity within the AIF–CHCHD4 axis, indicating that productive interactions and mitochondrial functions can be maintained across distinct AIF conformational states. These results refine current models of AIF activation and support a more dynamic framework linking redox sensing, dimerization and mitochondrial protein homeostasis.

## Results

### Dimerization of AIF conditions its mitochondrial homeostasis role

To investigate the mitochondrial and cellular consequences of AIF deficiency or impaired dimerization, mouse embryonic fibroblasts (MEFs) expressing different human AIF variants were generated. Thus, a stable AIF Knockout cell line (AIF^KO^ MEFs), previously generated by Dr. Susin’s laboratory (Delavallée *et al*, 2020), was transduced with lentiviral vectors encoding either the wild-type human AIF (hAIF WT) or the dimerization defective E413A/R422A/R430A mutant hAIF (hAIF 3M variant). These transductions generated the MEF-derived cell lines AIF^WT^ and AIF^3M^, respectively. (Fig. 2A). As a transduction control, AIF^KO^ MEFs were transfected with the same lentiviral plasmid containing GFP cDNA, generating the cell line AIF^KO^-GFP (hereafter AIF^KO^). After selection of the transduced cell lines, AIF expression and mitochondrial localization were evaluated by immunocytochemistry analysis. As shown in Fig. 2B, both hAIF^WT^ and hAIF^3M^ cell lines expressed the protein, which were properly imported and localized within mitochondria.

**Figure 2.**
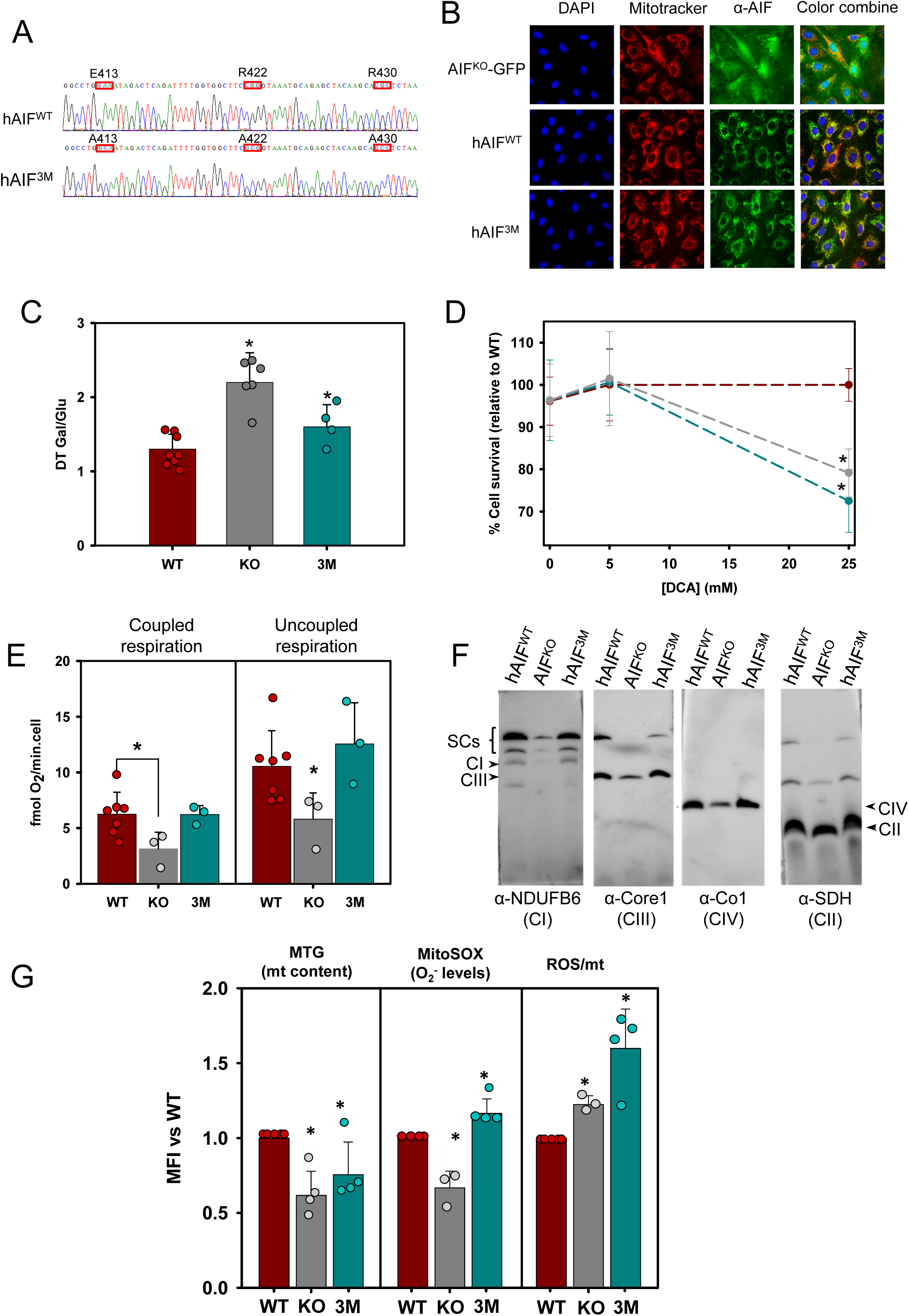
– Effect of AIF depletion and E413A/R422A/430A mutation on OXPHOS performance. **A** Chromatogram showing A1423C, C1449G&G1450C, and C1473G&G1474C transitions in human *AIFM1* gene leading to the E413A/R422A/R430A protein mutations. **B** Immunocytochemistry showing mitochondrial colocalization of hAIF in AIF^KO^ MEFS overexpressing GFP, or WT and 3M hAIF variants. From left to right: DAPI, AIF antibody, MitoTracker^TM^, and merged image (DAPI, blue; AIF, green; mitochondria, red). Note that green fluorescence observed in AIF^KO^ is due to GFP and does not represent AIF detection. **C** Growth ratio for hAIF^WT^, AIF^KO^ and hAIF^3M^ cell lines in medium containing galactose versus glucose-rich medium. *(P*=0.002 by non-parametric Kruskal– Wallis test; pairwise comparisons by Mann–Whitney *U* test: *P*=0.0019 for hAIF^WT^ vs. AIF^KO^, and *P*=0.0415 for hAIF^WT^ vs. hAIF^3M^). Data are expressed as mean ± SD of the mean. Sample sizes are as follows: n=8, 6 and 4 biological replicates for hAIF^WT^, AIF^KO^ and hAIF^3M^, respectively. **D** Cell viability of hAIF^WT^ (dark red), AIF^KO^ (dark grey) and hAIF^3M^ (dark cyan) cell lines after 72 h of 5 or 25 mM DCA treatment, measured by MTT assay and normalized to untreated hAIF^WT^ cells. Data are shown as mean ± SD (n = 6 biological replicates). Asterisks indicate statistical significance (\**P* < 0.05). Overall significance: *P*=0.0019 by non-parametric Kruskal–Wallis test; pairwise comparisons by Mann–Whitney *U* test: *P*=0.0039 for hAIF^WT^ vs. AIF^KO^, and hAIF^WT^ vs. hAIF^3M^. **E** Oxygen consumption rate in intact cells under basal conditions (coupled respiration, left) and after addition of the uncoupler DNP (uncoupled respiration, right). Data are presented as mean ± SD (n = 7, 3 and 3 biological replicates for hAIF^WT^, AIF^KO^ and hAIF^3M^, respectively). Asterisks indicate statistical significance (\**P* < 0.05). Overall significance by non-parametric Kruskal Wallis test: *P* = 0.0877 for coupled and *P* = 0.0292 for uncoupled respiration; pairwise comparisons by Mann–Whitney *U* test: *P*=0.0527 and *P* = 0.0167 for hAIF^WT^ vs. AIF^KO^ under coupled and uncoupled respiration, respectively. **F** Immunoblot of assembled supercomplexes in digitonin-permeabilized mitochondria separated by BNGE and probed with specific antibodies for CI (anti-NDUFB6), CIII (anti-Core1), CIV (anti-Co1) and CII (anti-SDHA). **G** Mitochondrial content, ROS production and ROS/mitochondrial content ratio measured by flow cytometry after staining with MitoTracker^TM^ green and MitoSOX^TM^. Data are represented as mean ± SD (n = 5, 4 and 5 biological replicates for mitochondrial content; n = 5, 3 and 4 biological replicates for ROS measurements). Asterisks indicate statistical significance (*P* < 0.05). Overall significance by non-parametric Kruskal Wallis test: *P* = 0.0463 for mitochondrial content, *P* = 0.0079 for mitochondrial superoxide production and *P* = 0.1237 for mtROS normalized by mt content; pairwise comparisons by Mann–Whitney *U* test: *P*=0.0143 and *P* = 0.0253 for hAIF^WT^ vs. AIF^KO^ for mitochondrial content and ROS production, respectively; and *P*=0.0143 hAIF^WT^ vs. hAIF^3M^ for mitochondrial ROS production.

To determine the relevance of AIF dimerization in mitochondrial homeostasis, we compared the effects of AIF loss and impaired dimerization on MEFs growth under the following metabolic conditions: glucose-containing medium and medium in which glucose was replaced by galactose, thereby forcing cells to rely on OXPHOS rather than fermentation for energy production. Whereas all cell lines showed similar growth rates in glucose-containing medium, both AIF^KO^ and hAIF^3M^ cells exhibited a reduced growth in galactose enriched-medium compared to hAIF^WT^ cells. Consequently, the galactose-to-glucose doubling time ratio was significantly higher in both cell lines, consistent with OXPHOS impairment (Fig. 2C).

Similarly, forcing OXPHOS through treatment with dichloroacetate (DCA), an inhibitor of pyruvate dehydrogenase kinase, significantly impaired cell growth in both AIF^KO^ and hAIF^3M^ cell lines (Fig. 2D). In both cases, DCA inhibition and galactose medium, the reduction in growth was less pronounced in hAIF^3M^ cells than in the AIF^KO^ cell line, suggesting a more severe impairment in OXPHOS in the absence of AIF.

To further address the effects of AIF variants on mitochondrial function, OXPHOS activity and organization were analyzed. As shown in Fig. 2E-F, the absence of AIF promoted a significant decrease in mitochondrial respiration both as coupled oxygen consumption and as uncoupled respiration, together with a strong reduction in the levels of mitochondrial respiratory complexes and supercomplexes, as previously described (Delavallée *et al*., 2020; Modjtahedi *et al*, 2015; Vahsen *et al*, 2004). Since AIF participates in the import and oxidative folding of specific OXPHOS subunits (Salscheider *et al*., 2022; Vahsen *et al*., 2004), its absence likely causes a severe impairment of these processes, contributing to mitochondrial dysfunction.

Consistent with this, AIF^KO^ cells displayed a significant reduction in mitochondrial content (Fig 2G) and lower mitochondrial ROS levels, measured as superoxide, than hAIF^WT^ cells (Fig. 2G). However, normalization of ROS levels to mitochondrial mass revealed a significant increase in superoxide production in AIF^KO^ cells (∼ 23% increase), indicating enhanced mitochondrial oxidative stress, despite the overall reduction in mitochondrial content.

In contrast, mitochondrial respiratory capacity in hAIF^3M^ cells was comparable to that of hAIF^WT^ cells (Fig. 2E). However, the organization of the respiratory chain was altered. Specifically, the levels of complexes I and III incorporated into supercomplexes were reduced relative to their free forms, reaching approximately 90% and 70% of those in hAIF^WT^ cells, respectively (mean of three biological replicates; Fig. 2F). Furthermore, despite a reduction in mitochondrial content similar to that observed in AIF^KO^ cells (Fig. 2G), hAIF^3M^ cells exhibited increased mitochondrial ROS production (Fig. 2G). This phenotype became even more pronounced after normalization to mitochondrial content, showing an approximately 60% increase compared with hAIF^WT^ cells, which is consistent with impaired OXPHOS performance.

These findings suggest that impaired AIF dimerization partially compromises its mitochondrial function rather than abolishing it, consistent with a model in which distinct AIF conformational states contribute differentially to mitochondrial homeostasis.

### Dimerization of AIF modulates the formation and composition of the mitochondrial AIF:CHCHD4 complex

CHCHD4 interacts with AIF during mitochondrial import and oxidative folding, and recent structural studies indicate that this interaction is optimized in the NADH-stabilized dimeric state (Brosey *et al*., 2025; Hangen *et al*., 2015; Rothemann *et al*., 2025). Based on these observations, we investigated the impact of AIF dimerization on CHCHD4 expression and mitochondrial import.

To properly interpret the effects of impaired AIF dimerization on CHCHD4, AIF expression was first evaluated in the different MEF-derived cell lines. As expected, AIF expression was undetectable in AIF^KO^ cells at both the mRNA and protein levels (Fig. 3A, B). In contrast, hAIF^3M^ cells showed higher AIF transcript levels (Fig. 3A) but significantly reduced protein abundance in whole-cell homogenates compared with hAIF^WT^ cells (Fig. 3B), suggesting decreased stability of the mutant protein.

**Figure 3.**
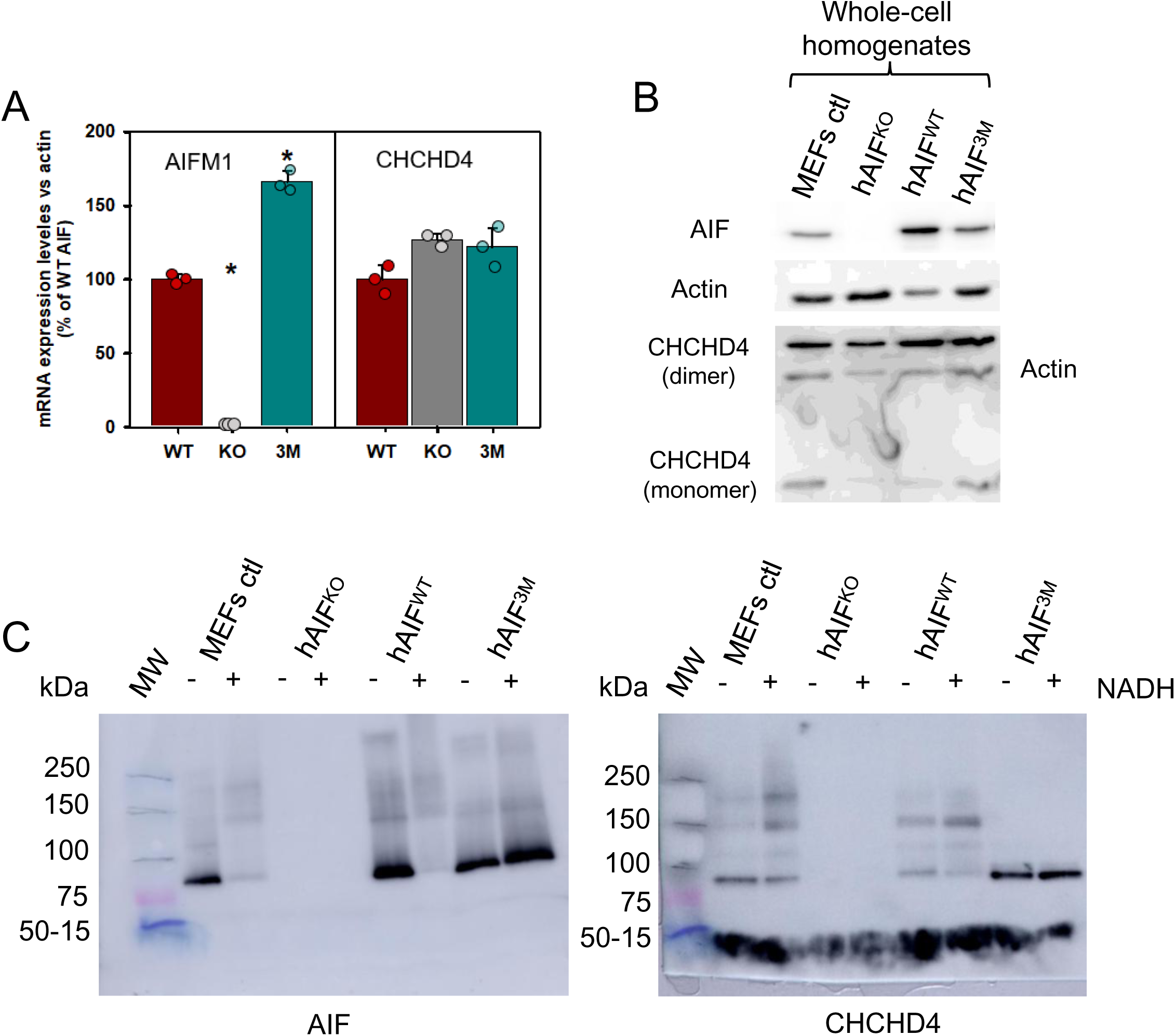
– The role of AIF dimerization in its interaction with CHCHD4. **A** qPCR analysis of the expression of *AIF* and *Chchd4* mRNAs relative to actin in AIF^WT^, AIF^KO^ and AIF^3M^ cells. Data are presented as mean ± SD (n = 3 technical replicates). Asterisks indicate statistical significance (*P* < 0.05). Overall significance by non-parametric Kruskal Wallis test: *P* = 0.0273 for *AIF* relative expression; pairwise comparisons by Mann–Whitney *U* test: *P*=0.0495 for hAIF^WT^ vs. AIF^KO^ and for hAIF^WT^ and hAIF^3M^. **B** Western blot analysis of AIF and CHCHD4 proteins in whole-cell protein extracts, normalized to actin and upon separation by SDS-PAGE. **C** Western blot of AIF:CHCHD4 interaction in digitonin-permeabilized mitochondria from hAIF^WT^, AIF^KO^-GFP, hAIF^3M^ and control MEFs cell lines, separated by CN-PAGE. The blots were probed with anti-AIF (left) and anti-CHCHD4 (right) specific antibodies.

Neither AIF depletion nor expression of the hAIF^3M^ variant affected CHCHD4 mRNA levels (Fig. 3A, right panel). However, total CHCHD4 protein abundance was reduced in both AIF^KO^ and hAIF^3M^ whole-cell homogenates (Fig. 3B), supporting the previously described posttranscriptional role of AIF in CHCHD4 stability and/or mitochondrial import (Hangen *et al*., 2015; Meyer *et al*., 2015).

To assess the functional relevance of AIF dimerization, mitochondrial extracts were analyzed by CN-PAGE under basal conditions and after NADH incubation. As expected, AIF^WT^ (both the endogenous murine form in control MEFs and the human version in hAIF^WT^ cells) shifted from a monomer-dimer equilibrium toward a predominantly dimeric state upon NADH-induced reduction, whereas hAIF^3M^ remained monomeric under both conditions (Fig. 3C, left panel). Moreover, CHCHD4 was detected both in complex with AIF (exhibiting higher affinity for the dimeric form of AIF) and in a free form, independently of its redox state. Indeed, in control MEFs and hAIF^WT^ cells the ratio of CHCHD4 bound to AIF increased approximately 2– and 6-fold in the presence of NADH, respectively. In AIF^KO^ cells, CHCHD4 was still imported into mitochondria but was detected exclusively in its free form (Fig. 3C, right panel).

### Dimerization defects in AIF compromise redox function and NADH-linked structural organization

To better define the contribution of AIF dimerization to its mitochondrial function, we performed a comparative molecular characterization of WT_Δ77_ and two variants, 3M_Δ77_ and H454A_Δ77_, which disrupt NADH-linked quaternary assembly at different stages. The 3M_Δ77_ variant is unable to stabilize the dimeric conformation upon flavin reduction by NADH (Ferreira *et al*., 2014), and only releases the regulatory C-loop in the presence of a large excess of NADH (SI appendix). In contrast, the H454A_Δ77_ variant, which substitutes a key active-site residue required for CTC stabilization, adopts a dimer-permissive conformation while retaining an intact C-loop in both oxidized and reduced states (Brosey *et al*., 2016) (SI appendix). Notably, these AIF_Δ77_ variants lack the membrane-anchoring helix but retain the IMS soluble portion, thus representing physiologically relevant mitochondrial isoforms.

Purified proteins were analyzed by BS3 cross-linking followed by SDS-PAGE allowing discrimination between monomeric and dimeric conformations (Villanueva *et al*, 2019) (Fig. S1A). Consistent with its reported defect in NADH-induced assembly, 3M_Δ77_ remained exclusively monomeric regardless of redox conditions. In contrast, H454A_Δ77_ formed cross-linked dimers both in the presence and absence of NADH, indicating preservation of dimer-compatible conformations despite defective CTC stabilization. Gel filtration chromatography further revealed that 3M_Δ77_ consistently eluted as a monomer (^app^MW of 62 kDa), whereas WT_Δ77_ predominantly adopted a dimeric conformation (∼170 kDa) in the presence of NADH (SI appendix and Fig. S1B-C and S1E-F). Conversely, the H454A_Δ77_ variant exhibited two dimer conformations (^app^MW of ∼115 and ∼150 kDa) both in the presence and absence of NADH (SI appendix and Fig. S1G-H). Importantly, the 3M_Δ77_ and H454A_Δ77_ variants exhibited spectroscopic properties similar to those of WT_Δ77_ (Villanueva *et al*., 2019), indicating that these mutations did not substantially perturb flavin binding or global protein folding (SI appendix and Fig. S2A-B). Together, these data establish that the mutations selectively alter NADH-linked structural organization without globally destabilizing the protein.

AIF exhibits a NADH oxidase activity in mitochondria, which can be studied *in vitro* using the steady-state DCPIP-dependent diaphorase assay. As shown in Table 1 and Fig. S2C, the 3M_Δ77_ and H454A_Δ77_ variants showed reduced NADH affinity (by ∼8– and ∼4-fold, respectively) but similar or slightly higher turnover rates compared to the WT_Δ77_. Consequently, 3M_Δ77_ and H454A_Δ77_ variants were, respectively, 14– and 2-fold less efficient at oxidizing NADH than WT_Δ77._

**Table 1.** Steady-state and pre-steady state kinetic parameters of AIF_Δ77_ variants.

| Sample | Steady-state |  |  | Pre-steady-state |  |  | CTC |
| --- | --- | --- | --- | --- | --- | --- | --- |
| | $k_{cat}$<br>(s<sup>-1</sup>) | $K_m^{NADH}$<br>(μM) | $k_{cat}/K_m^{NADH}$<br>(s<sup>-1</sup> mM<sup>-1</sup>) | $k_{HT}$<br>(s<sup>-1</sup>) | $K_d^{NADH}$<br>(μM) | $k_{HT}/K_d^{NADH}$<br>(s<sup>-1</sup> mM<sup>-1</sup>) | Half-life<br>(min) |
| **WT** | 2 ± 0.1 | 50 ± 9 | 41 ± 8 | 2.3 ± 0.3 | 1544 ± 83 | 1.5 ± 0.1 | 20 |
| **3M** | 1.1 ± 0.1 | 401 ± 69 | 2.9 ± 0.4 | 1.2 ± 0.1 | 1569 ± 253 | 0.8 ± 0.1 | ND |
| **H454A** | 4.7 ± 0.5 | 194 ± 6 | 26 ± 3 | 4.7 ± 0.5 | 243 ± 36 | 20 ± 5 | NS |
| **R422Q** | 1.6 ± 0.2 | 360 ± 50 | 5 ± 1 | 2.2 ± 0.7 | 1733 ± 160 | 1.3 ± 0.5 | 5 |
| **WT:N-CHCHD4** | 1.2 ± 0.2 | 16 ± 5 | 75 ± 7 | 135 ± 34 | 320 ± 30 | 417 ± 68 | 52 |
| **3M:N-CHCHD4** | 1.2 ± 0.1 | 10 ± 2 | 115 ± 11 | 60 ± 0.7 | 135 ± 20 | 449 ± 70 | 24 |
| **H454A:N-CHCHD4** | 6.8 ± 0.9 | 20 ± 2 | 350 ± 90 | 242 ± 17<sup>a</sup> | 406 ± 80 | 603 ± 78 | 0.6 |
| **R422Q:N-CHCHD4** | 1.3 ± 0.1 | 80 ± 11 | 18 ± 4 | ND |  |  | 84 |
| **WT:CHCHD4** | 1.1 ± 0.1 | 15 ± 3 | 74 ± 15 | $k_{HT} \sim 0.9$, [NADH] independent | | | ≥ 1440 |
| **3M:CHCHD4** | 1.6 ± 0.1 | 19 ± 6 | 85 ± 23 | 1.7 ± 0.1 | 1696 ± 14 | 1.0 ± 0.1 | 27 |
| **H454A:CHCHD4** | 5.5 ± 0.4 | 22 ± 2 | 255 ± 5 | 205 ± 4<sup>a</sup> | 385 ± 4 | 533 ± 16 | 2 |

Assays were performed at 25 °C in 50 mM potassium phosphate, pH 7.4. (n=3, mean ±SD); ND: not determined under assay conditions; NS: not stabilized.

To evaluate the impact of mutations on hydride transfer (HT) from NADH to FAD cofactor, we used stopped-flow transient kinetic analyses. A kinetic comparison between WT_Δ77_ and the apoptotic WT_Δ101_ isoform is additionally provided in the SI appendix. In 3M_Δ77_, full FAD reduction was accompanied by the progressive formation of FADH^-^:NAD^+^ CTC with an intensity similar to that in the WT_Δ77_ (Fig. 4A-B), indicating that the mutation does not abolish CTC formation *per se*. However, unlike the WT_Δ77_, which had a CTC half-life time of 20 minutes, the 3M_Δ77_ could not be fully reduced at stoichiometric concentrations of coenzyme, thereby hindering CTC accumulation and determination of its reactivity towards O_2_. In addition, HT rates were modestly decreased, making 3M_Δ77_ ∼2-fold less efficient than WT_Δ77_ in oxidizing the coenzyme (Table 1). In contrast, H454A_Δ77_ variant displayed a slightly faster HT (∼2-fold higher) together with higher NADH affinity (6-fold lower *K*_d_), enhancing HT efficiency ∼13-fold (Table 1). Consistent with this behavior, H454A_Δ77_ failed to stabilize a detectable CTC in the presence of NADH (Fig. 4C), in agreement with previous reports on H454 variants (Brosey *et al*., 2016; Churbanova & Sevrioukova, 2008; Villanueva *et al*., 2015). Thus, while dimerization defects primarily impair stabilization of NADH-dependent redox states, disruption of the H454-mediated pathway uncouples HT from productive CTC formation. Altogether, these findings demonstrate that the dimer-interface integrity and the C-loop conformational dynamics are essential for coupling AIF’s redox activity with NADH-induced structural transitions during dimer formation.

**Figure 4.**
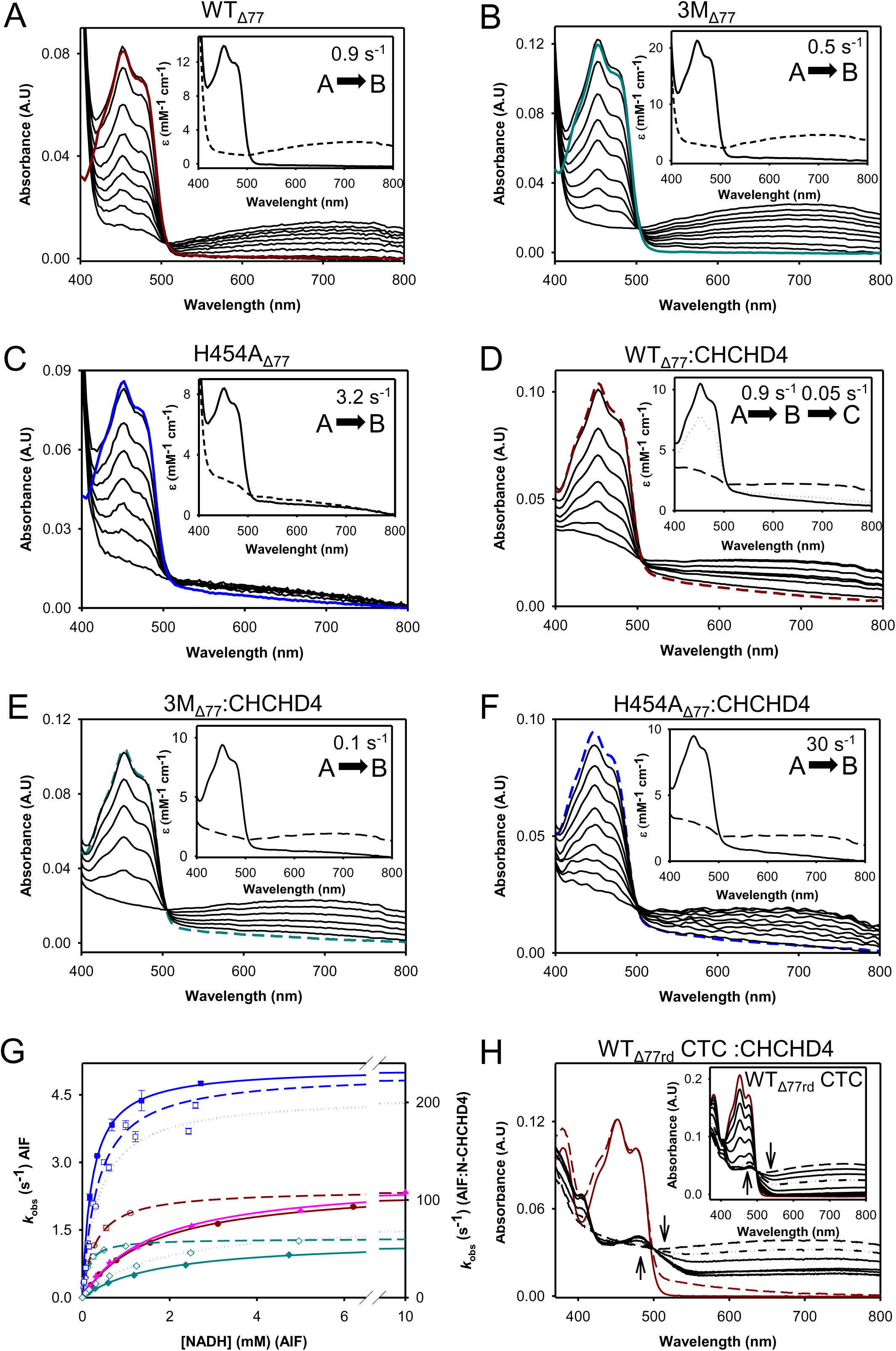
– CHCHD4 modulates AIF redox properties. **A-F** Time course of HT from NADH to AIF_Δ77_ variants. Spectral evolution for flavin reduction of (A) WT_Δ77_, (B) 3M_Δ77_, (C) H454A_Δ77_ and (D) WT_Δ77_:CHCHD4, (E) 3M _Δ77_:CHCHD4 and (F) H454A_Δ77_:CHCHD4. Insets show simulated spectral species from global fitting either to a single step (A→B) or two steps (A→B→C) kinetic model. Initial spectra of WT_Δ77,_ 3M_Δ77_ and H454A_Δ77_ before NADH mixing, without or with CHCHD4, are shown as solid or dashed dark red, dark cyan and blue lines, respectively. Spectral changes were acquired at 0.05, 0.25, 0.55, 0.9, 1.25, 1.7, 2.3, 3.3 and 19.5 s for WT_Δ77_, at 0.18, 0.88, 1.6, 2.28, 2.98, 4.9, 6.65, 10.15 and 20 s for 3M_Δ77_, at 0.003, 0.033, 0.078, 0.18, 0,33, 0.7 and 2 s for H454A_Δ77_, at 0.25, 2.5, 7.25, 14.75, 24.75, 49.75 and 100 s for WT_Δ77_:CHCHD4, at 0.08, 1.52, 4, 8.2, 17.5 and 80 s for 3M _Δ77_:CHCHD4 and at 0.001, 0.008, 0.014, 0.022, 0.029, 0.043, 0.058, 0.087 and 0.4 s for H454A_Δ77_:CHCHD4. **G** NADH concentration dependence of the observed rate constants for flavin reduction of WT _Δ77_ (dark red closed circles), 3M_Δ77_ (dark cyan closed diamonds) and H454A_Δ77_ (blue closed squares) variants and their corresponding complexes with CHCHD4 or its N-terminal peptide (open symbols using the above colour code). Solid, dash and dotted lines represent the fits of experimental data to Equation 4 respectively for the AIF_Δ77_ variants, and the corresponding AIF_Δ77_: N-CHCHD4 and AIF_Δ77_:CHCHD4 complexes. **H** Spectral evolution of the reactivity of the WT_Δ77rd_ CTC towards O_2_ in the absence (inset) and presence of CHCHD4. CTC samples were prepared by mixing WT_Δ77_ with 1.5–fold excess NADH. Dark red solid and dashed lines represent the spectra of WT_Δ77rd_ in the absence or presence of CHCHD4, respectively, before mixing with NADH. Black dashed dotted lines represent the WT_Δ77_ CTC half-life time.

### CHCHD4 binding significantly modulates AIF redox properties and conformational dynamics, functionally compensating the dimerization defect

To determine the relevance of AIF dimerization and C-loop dynamics on CHCHD4 interaction, we first characterized the binding of the AIF_Δ77_ variants to the full-length CHCHD4 or to a peptide corresponding to its N-terminal 27 residues (herein N-CHCHD4), using isothermal titration calorimetry (ITC) (Fig. EV1, Table EV1). In contrast to the apoptotic WT_Δ101ox_ (Brosey *et al*., 2025; Romero-Tamayo *et al*., 2021; Rothemann *et al*., 2025), the mitochondrial protein forms a stable and detectable complex with CHCHD4. Both WT_Δ77ox_ and the dimer-permissive H454A_Δ77ox,_ mutant, which retains a surface-bound C-loop, interacted with CHCHD4 through enthalpically driven binding accompanied by unfavorable entropic contributions, consistent with specific complex formation. Binding affinity was stronger for the H454A_Δ77ox_ variant than for WT_Δ77ox_ (*K*_d_ ∼0.9 µM and ∼7.7 µM, respectively), with values comparable to those reported for murine WT_Δ101_ (Fagnani *et al*., 2024). Moreover, H454A_Δ77ox_ interacted with N-CHCHD4 with a thermodynamic profile similar to that observed with the full-length chaperone, although exhibiting a stronger enthalpic contribution (–20 vs –12 kcal/mol). However, affinity remained higher for full-length CHCHD4, indicating that additional contacts beyond the N-terminal region contribute to the complex stabilization. These observations indicate that the C-loop displacement is not strictly required for CHCHD4 engagement, although it likely contributes to interaction optimization.

In the presence of an excess of NADH, interaction with both full-length CHCHD4 and N-CHCHD4 was detected in all variants. WT_Δ77rd_ and 3M_Δ77rd_ exhibited similar thermodynamic profiles, characterized by favourable enthalpic contributions, consistent with previous observation for the apoptotic WT_Δ101rd_ (Romero-Tamayo *et al*., 2021). Reduction increased the affinity of WT_Δ77_ for CHCHD4 by approximately 10-fold. Although 3M_Δ77rd_ remained competent for interaction, both enthalpic and entropic contributions were reduced (ΔH ∼6 kcal/mol and –TΔS ∼5 kcal/mol lower, respectively), indicating weakened interaction. Thus, disruption of the dimer interface weakens but does not abolish CHCHD4 binding.

Interestingly, H454A_Δ77rd_ exhibited stronger binding to both CHCHD4 and N-CHCHD4 (∼5-fold lower than in the oxidized state), accompanied by altered entropic contributions which became slightly favorable for CHCHD4 binding (–2.5 kcal/mol) but remained unfavorable for the peptide (6.9 kcal/mol). In addition, the enthalpic contribution to N-CHCHD4 binding was reduced compared to the oxidized state. Across all conditions and variants, ITC data supported a 1:1 binding stoichiometry between AIF and CHCHD4 or N-CHCHD4, in agreement with previous reports for WT_Δ101rd_ (Brosey *et al*., 2025; Hangen *et al*., 2015; Romero-Tamayo *et al*., 2021). Binding data to the N-terminal peptide further confirm that the interaction is independent of the redox-active CHCHD4 CPC motif and primarily mediated by structural determinants within its N-terminus (Brosey *et al*., 2025).

To determine whether CHCHD4 binding influences AIF conformational stability, we next analyzed the thermal stability of AIF_Δ77_ variants in the presence of full-length CHCHD4 and N-CHCHD4 (Fig. S3, Table S1). No significant changes were observed for WT_Δ77_ or 3M_Δ77_ in either redox state. In contrast, both CHCHD4 and N-CHCHD4 reduced the thermal stability of H454A_Δ77rd_, resembling the behavior of the reduced forms of WT_Δ77_ and 3M_Δ77_ alone, previously attributed to increased C-loop flexibility and openness (Romero-Tamayo *et al*., 2021; Villanueva *et al*., 2019). This effect suggests that CHCHD4 binding promotes conformational rearrangements in AIF associated with increased C-loop accessibility.

Size-exclusion chromatography further confirmed formation of 1:1 AIF:CHCHD4 complexes for all AIF variants independently of their redox state (SI appendix and Fig. S1B-H). Notably, a significant fraction of 3M_Δ77rd_ remained unbound, indicating reduced complex stability. In addition, CHCHD4 promoted the stabilization of higher-order assemblies in all variants except 3M_Δ77ox_. Similar assemblies were observed in mitochondrial extracts (Fig. 3C) as well as in *in vitro* cross-linking assays for mAIF_Δ101_:CHCHD4 mixtures (Fagnani *et al*., 2024), suggesting that CHCHD4 influences AIF oligomerization. Together, these data indicate that CHCHD4 interaction dynamically remodels AIF conformational and oligomeric states.

We next evaluated whether CHCHD4 binding affects AIF redox properties (Table 1, Fig. EV2). Under steady-state conditions, both CHCHD4 and N-CHCHD4 enhanced coenzyme affinity across all variants (∼3, ∼20 and ∼9-fold for WT_Δ77_, 3M_Δ77_ and H454A_Δ77_, respectively), shifting apparent affinities toward physiologically relevant NADH concentrations (Rothemann *et al*., 2025). Consequently, mutant proteins recovered affinity values close to those of WT_Δ77_. Notably, catalytic efficiency in 3M_Δ77_ increased ∼38-fold in the presence of CHCHD4, reaching values comparable to those for WT_Δ77_. Thus, CHCHD4 interaction can partially compensate for defects associated with impaired dimer stabilization.

HT was also strongly enhanced by both CHCHD4 and N-CHCHD4 (Table 1, Figs 4D-G and EV2A-C). In particular, the N-terminal peptide accelerated HT across all variants (∼400, ∼450 and ∼30 fold for WT_Δ77_, 3M_Δ77_ and H454A_Δ77,_ respectively), rendering the reductive half-reaction extraordinarily efficient. These effects occurred without detectable changes in AIF redox potential (–358 mV for both WT_Δ77_ alone and WT_Δ77_:N-terminal complex), supporting a mechanism based on the remodeling of catalytic geometry rather than the alteration of intrinsic redox thermodynamics. In contrast, no effects were observed for 3M_Δ77_ in the presence of the full-length chaperone (Fig. 4E), while the HT reaction of WT_Δ77_ turned out NADH-independent (k_HT_ ∼0.9, similar to *k*_cat_ under steady-state conditions). H454A_Δ77_ maintained HT reaction parameters similar to those observed with the N-CHCHD4. Remarkably, the binding of CHCHD4 or its N-terminal peptide induced spectral changes in the flavin environment (Fig. 4H, Fig. EV2D), consistent with the proposed structural remodeling toward a catalytically competent conformation.

N-CHCHD4 additionally increased CTC stability in WT_Δ77_ and 3M_Δ77_ (Table 1, Fig. EV2D-E (inset), increasing its lifetime in WT_Δ77_ (∼3-fold) and enabling its estimation in 3M_Δ77_ (24 min). Full-length CHCHD4 had a comparable effect on 3M_Δ77_ and formed a long-lived complex with WT_Δ77_ that persisted until protein destabilization (Fig. 4H). Remarkably, H454A_Δ77_ acquired detectable CTC features (Fig. EV2F) in the presence of either N-CHCHD4 or CHCHD4, with lifetimes of 0.6 and 2 min, respectively. Thus, CHCHD4 interaction promotes conformational states compatible with CTC stabilization even in variants defective in the canonical NADH-dependent allosteric pathway.

Collectively, these findings show that CHCHD4 binding coordinately remodels AIF conformational and redox properties, enhancing catalytic performance and partially restoring function in dimerization– or redox-defective variants.

### The N-terminal peptide of CHCHD4 partially restores redox properties and thermal stabilization in the pathogenic AIF R422Q variant

Given the positive effects of N-CHCHD4 on our engineered AIF mutants, we next explored whether this peptide could partially rescue the defective redox properties of a pathogenic AIF mutant with impaired dimer stabilization. For this purpose, we selected the disease-associated R422Q variant, previously reported to reduce dimeric AIF levels in patient-derived neurons (Qiu *et al*., 2023).

Following purification, BS^3^ cross-linking assays confirmed that R422Q_Δ77_ retained the ability to dimerize in the presence of NADH (Fig. 5A). Moreover, R422Q_Δ77_ displayed spectroscopic properties comparable to those of WT_Δ77_, indicating that the mutation did not affect the flavin environment or the overall protein conformation (Fig. S2A-B). However, the R422Q_Δ77rd_ exhibited a greater conformational destabilization compared with WT_Δ77_ (Fig. S3A and Table S1), along with decreased coenzyme affinity and severely impaired CTC stabilization (Fig. 5B inset and Table 1). Notably, the HT reaction remained unaffected (Table 1), although the intensity of the CTC band was slightly lower compared with WT_Δ77_ (Fig. 5C). In summary, these results suggest that the R422Q mutation induces destabilization of the AIF dimer by weakening the CTC stabilization and reducing the NADH affinity. In the presence of N-CHCHD4, the conformational stability of the R422Q_Δ77rd_ minimally improved (Fig. S3B, Table S1). However, it partially restored the R422Q_Δ77_ redox function, significantly enhancing CTC stabilization and moderately improving coenzyme affinity (CTC lifetime ∼16 fold greater and *K*_d_ ∼5 fold lower compared to R422Q_Δ77_ alone) (Fig. 5B, Figure S2C and Table 1). These effects were accompanied by spectral changes in the flavin environment consistent with active-site remodeling (Fig. 5B). Thus, interaction with the CHCHD4 N-terminal region partially restores functional redox properties despite persistent destabilization of the mutant conformation.

**Figure 5.**
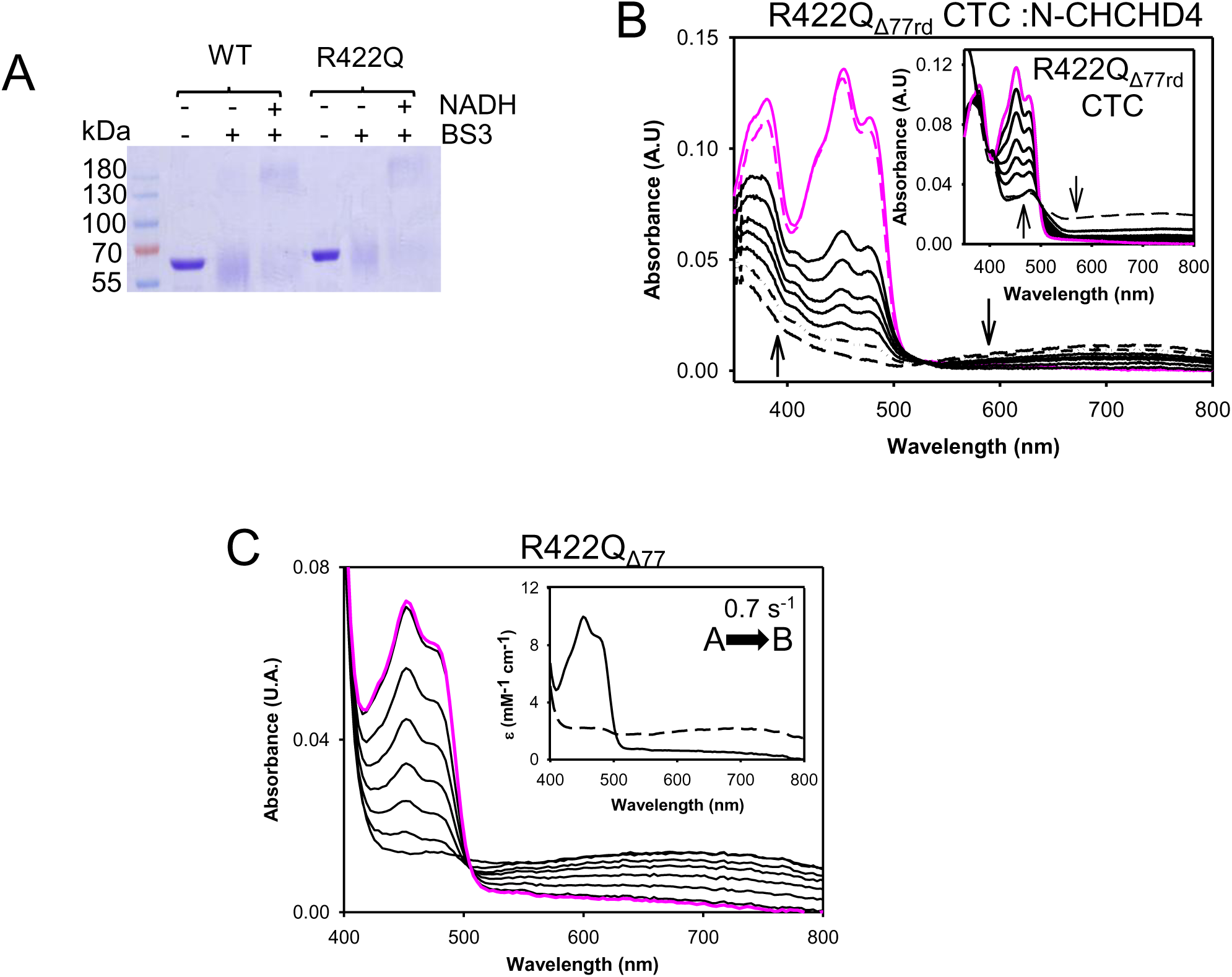
– Impact of the R422Q_Δ77_ mutation in AIF redox properties. **A** Chemical cross-linking of R422Q_Δ77_ using a 100-fold excess of the BS^3^ cross-linker without or with 100-fold excess of NADH. After 30 minutes of incubation, reactions were quenched with bromophenol-containing sample buffer and resolved by 12% SDS-PAGE. **B** Spectral evolution of the reactivity of the R422Q_Δ77rd_:NAD^+^ CTC towards O_2_ in the absence (inset) and presence of the N-CHCHD4. CTC samples were prepared by mixing R422Q_Δ77ox_ with 1.5–fold excess of NADH. Pink solid and dashed lines correspond to the spectra of R422Q_Δ77ox_ in the absence or presence of CHCHD4, respectively, before mixing with NADH. Black dashed dotted lines correspond to WT_Δ77_ CTC half-life time. **C** Time course of HT from NADH to R422Q_Δ77_. Spectral evolution of flavin reduction in the R422Q_Δ77_ variant. The spectrum of R422Q_Δ77_ prior to mixing with NADH is shown as a solid pink line. The inset shows the simulated spectral species predicted by global fitting of the experimental data to a single step (A→B) kinetic model. Spectral changes were recorded at 0.03, 0.33, 0.72, 1.2, 1.8, 2.8 and 15 s.

Overall, these results further support the concept that AIF remains functionally plastic and responsive to partner-induced allosteric modulation, even when primary dimer-stabilizing interactions are compromised.

## Discussion

A significant subset of inherited mitochondrial diseases is linked to defects in the IMS protein import machinery, a system responsible for the translocation and oxidative folding of hundreds of nuclear-encoded proteins essential for normal mitochondrial function (Wischhof *et al*., 2022). These substrates include components of the OXPHOS system, the cristae-remodeling MICOS complex, and mitochondrial import complexes (Erdogan & Riemer, 2017; Habich *et al*, 2019). The primary IMS import pathway is the DRS, which couples protein import to chaperone-assisted folding through the activity of CHCHD4, targeting a specific class of small soluble proteins containing cysteine motifs (CX_3_C and CX_9_C) and promoting their stabilization via disulfide bond formation (Al-Habib & Ashcroft, 2021). Within this framework, AIF has emerged as a critical regulator of IMS proteostasis by facilitating the CHCHD4 function.

Initial models proposed that AIF acts primarily as an import platform that facilitates CHCHD4 positioning and substrate engagement at the TOM complex (Brosey *et al*., 2025; Hangen *et al*., 2015; Meyer *et al*., 2015). Subsequent work further demonstrated that NADH-dependent AIF dimerization promotes conformational rearrangements that enhance CHCHD4 activity through exposure of its CPC catalytic motif (Brosey *et al*., 2025). Our findings extend this model by demonstrating that the NADH-stabilized AIF dimer represents a highly efficient, but not exclusive, functional state for CHCHD4 engagement. Rather than operating as a binary on/off switch, the AIF– CHCHD4 system behaves as a conformationally dynamic ensemble, in which distinct AIF redox and oligomeric states retain varying capacities to sustain CHCHD4 activity (Fig. 6).

**Figure 6.**
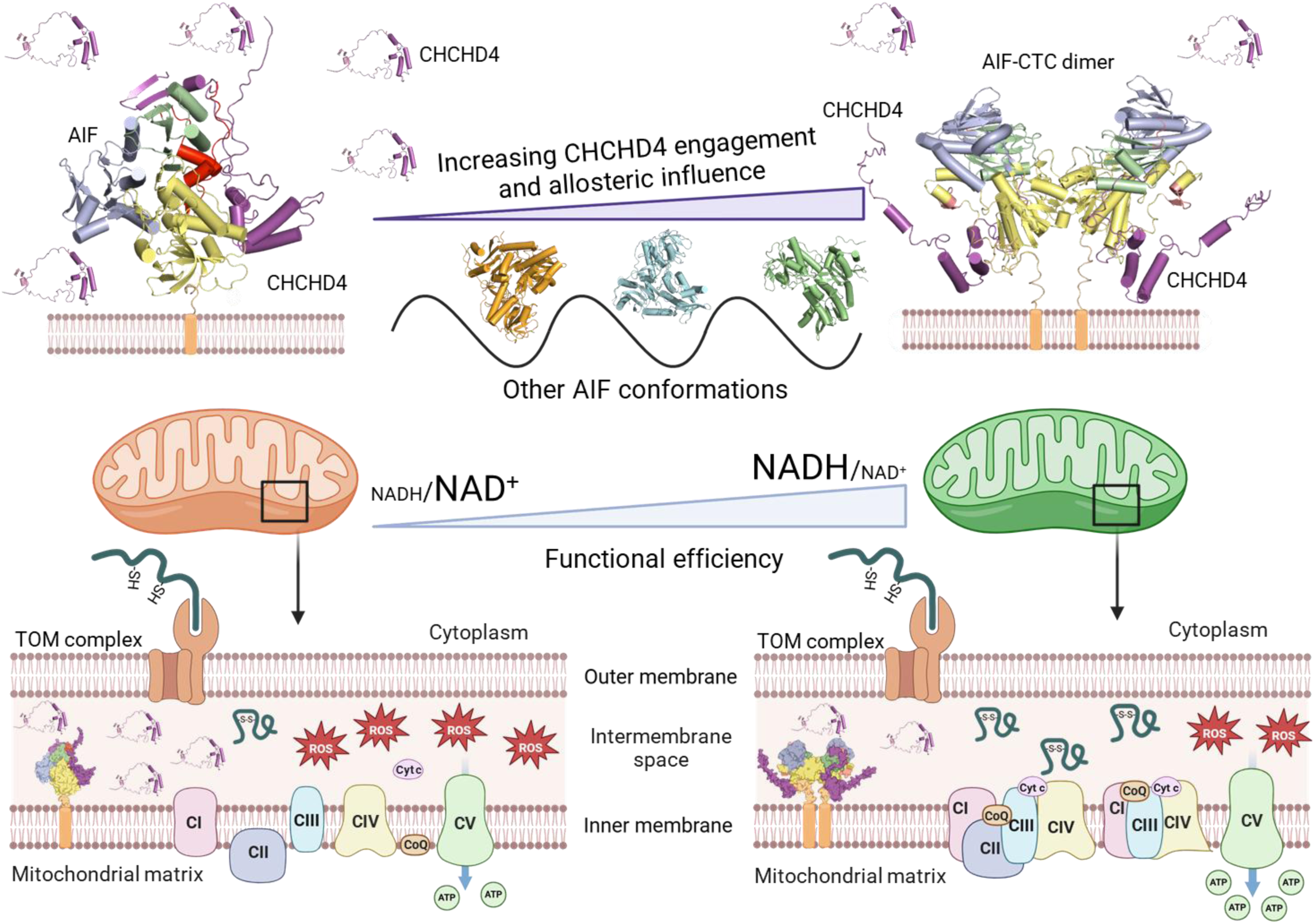
– Functional plasticity of AIF:CHCHD4 interaction: Under conditions of a low NADH/NAD^+^ ratio in the IMS, monomeric AIF possesses a functional but restricted capacity to bind and sustain CHCHD4 function, leading to protein import decrease through the DRS, OXPHOS deficiency and increased ROS levels. Upon an increase in NADH levels, AIF undergoes dimerization and establishes a more efficient complex with CHCHD4, exerting a bidirectional allosteric influence on the activities of both proteins. This long-lived, stable complex supports CHCHD4 activity in the DRS, OXPHOS performance, and mitochondrial homeostasis. The NADH-binding, FAD-binding and C-terminal domain, together with the C-loop and the β hairpin of AIF are coloured as previously described in Figure 1. CHCHD4 is displayed in magenta. The structural models were constructed using PDB entries 4B6V (AIF monomer), 9GQZ (dimeric AIF CTC in complex with CHCHD4), as well as the corresponding Alphafold-predicted complex reported by Pei et al. 2022. The figure was created with Biorender.

This concept is supported by our cellular analyses. Although hAIF^3M^ cells displayed impaired respiratory growth, reduced incorporation of respiratory complexes I and III into supercomplexes, increased mitochondrial ROS levels and elevated sensitivity to metabolic stress, these defects remained substantially milder than those observed in AIF^KO^ cells. Importantly, hAIF^3M^ cells retained substantial respiratory activity and maintained stable monomeric AIF:CHCHD4 complexes, supporting the model in which defective dimer stabilization reduces functional efficiency without fully abolishing AIF-dependent mitochondrial activity. In contrast, complete AIF deficiency caused a severe disorganization of the respiratory supercomplexes and a marked decline in OXPHOS performance (Hangen *et al*., 2015; Meyer *et al*., 2015). These graded phenotypes support a model in which monomeric or partially activated AIF conformations sustain residual CHCHD4 function, whereas the NADH-stabilized dimer defines the optimal state for supporting respiratory-chain biogenesis and oxidative folding.

Consistent with this idea, Mussulini et al. recently reported that elevated NADH levels in complex I-deficient cells strengthen the AIF–CHCHD4/MIA40 interaction and stabilize dimer-associated conformations that suppress AIF-dependent apoptosis by concealing its nuclear localization sequence (Mussulini *et al*, 2025). Although studied in a different functional context, these observations further reinforce the concept that NADH regulates the AIFM1–CHCHD4 interaction network and that distinct AIF conformational states may coordinate multiple mitochondrial outputs depending on the metabolic context.

Our data additionally indicate that AIF primarily regulates CHCHD4 functional organization and catalytic competence rather than its mitochondrial localization *per se*. Although CHCHD4 abundance was reduced in AIF^KO^ and hAIF^3M^ cells, this chaperone remained detectable and accumulated predominantly in a free/unbound mitochondrial pool. This observation aligns with reports in Harlequin mouse tissues and additional knockout cell models showing preserved mitochondrial CHCHD4 localization (Meyer *et al*., 2015; Salscheider *et al*., 2022; Wischhof *et al*, 2018), as well as with recent evidence indicating that direct AIF interaction is not required for CHCHD4 mitochondrial import (Brosey *et al*., 2025). Notably, while previous studies employing pre-fractionation or size-exclusion chromatography (SEC) prior to gel analyses primarily detected CHCHD4 bound to AIF (Salscheider *et al*., 2022), our native gel analyses reveal a persistent pool of unbound CHCHD4 along with the assembly-competent complex. These methodological differences reconcile apparent discrepancies in complex stoichiometry and further support a model in which AIF does not merely serve as an anchor, but dynamically regulates the conformational and oligomeric transitions required for productive DRS function.

The significance of this functional plasticity is further supported by our biochemical and biophysical analyses. CHCHD4 formed complexes with AIF at both redox states, although interaction affinity and stability were enhanced in the reduced dimeric conformations. Importantly, the dimerization-defective 3M variant displayed detectable CHCHD4 interaction, while the H454A_Δ77_ variant remained competent for partner binding despite lacking canonical NADH-dependent C-loop activation. These observations indicate that the NADH-induced C-loop displacement enhances, but is not a key allosteric prerequisite for, productive complex formation, thereby broadening current structural models centred on the fully activated dimeric state (Brosey *et al*., 2025; Rothemann *et al*., 2025). Instead, they support a more permissive interaction landscape in which CHCHD4 can also engage partially activated or non-canonical AIF conformations. Notably, this behaviour is consistent with structural observations previously reported (Fagnani *et al*., 2024), which capture CHCHD4 interaction with AIF in conformations that do not fully reflect the NADH-activated state, further supporting the existence of alternative binding-competent configurations.

In parallel, our results demonstrate that CHCHD4 binding actively remodels AIF redox chemistry in a bidirectional manner. Interaction with CHCHD4 or its N-terminal motif enhanced NADH affinity and CTC stabilization and stimulated HT kinetics, even in variants defective in dimer stabilization or redox coupling. During the preparation of this manuscript, Rothemann *et al* reported cryo-EM structures of AIF bound to Mia40 and AK2A, showing that partner binding engages the same C-terminal β-sheet in the reduced AIF dimer through β-strand complementation and compaction of the NAD-binding domain (Rothemann *et al*., 2025). Notably, this structural compaction, which reduces the distance between FAD and NAD, is fully consistent with our observations that CHCHD4 renders the HT reaction effectively independent of NADH concentration. Such partner-driven catalytic optimization may stabilize AIF activity and preserve DRS efficiency under fluctuating mitochondrial redox conditions.

Our biochemical analyses further indicate that this bidirectional allostery can functionally compensate for defective AIF conformations by restoring key redox properties associated with catalytic competence. In particular, CHCHD4 restored NADH affinity and prolonged CTC lifetime in the dimerization-defective 3M_Δ77_ variant, while partially recovering detectable CTC stabilization in H454A_Δ77_, despite disruption of the canonical H454-dependent allosteric pathway. These findings support a model in which partner-assisted stabilization of the C-terminal interaction platform can partially substitute for defective dimer-driven allosteric propagation.

Finally, the ability of the CHCHD4 N-terminal motif to partially restore redox function in the pathogenic R422Q_Δ77_ variant indicates that in disease-associated AIF conformations retain a degree of structural and functional tractability even when dimer stabilization is compromised. This framework provides a mechanistic basis for understanding how pathogenic AIFM1 variants impair mitochondrial homeostasis while preserving partial residual activity. Consistent with this view, we recently characterized a novel pathogenic variant (E336K) that similarly destabilizes the CTC and weakens CHCHD4 binding, yet retains residual mitochondrial function and amenability to molecular intervention (Ferrer *et al*., 2026). More broadly, our findings suggest that partner-assisted allosteric modulation may emerge as a tractable strategy to restore or enhance AIF function in mitochondrial diseases.

## Methods

### Experimental model and study participant details

*Escherichia coli* strains were grown in 2xYT medium at 37 °C.

All the eukaryotic cell lines were cultured at 37 °C in a 5% CO_2_ atmosphere in high glucose DMEM (Gibco) supplemented with 10% FBS (fetal bovine serum, Gibco). AIF^KO^ MEFs were obtained as described elsewhere (Delavallée *et al*., 2020). Cell lines expressing human AIF (hAIF^WT^) and its E413A/R422A/R430A variant (hAIF^3M^) were generated by lentiviral transduction of AIF^KO^ MEFs using the corresponding hAIF constructs. The negative control AIF^KO-GFP^ was obtained after GFP overexpression in AIF^KO^ MEFs. Transduced cell lines were isolated by growing the cell population in the presence of 800 µg/mL of G418 (Gibco).

### Lentiviral particles production and transduction

cDNAs encoding full length WT and E413A/R422A/R430A hAIF variants were generated by Mutagenex Inc. and subcloned into a lentiviral expression vector derived from pWPXLd (Tronolab), in which the GFP sequence was replaced by a neomycin resistance cassette (pWPXLd-IRES-Neo^R^). As a negative control, GFP cDNA was inserted into pWPXLd-IRES-Neo^R^ vector using the same restriction site (*Pme*I) as in the hAIF constructs to generate the AIF^KO-GFP^ cell line. Lentiviral particles carrying the hAIF^WT^-pWPXLd-IRES-Neo^R^, hAIF^3M^-pWPXLd-IRES-Neo^R^ or the GFP-pWPXLd-IRES-Neo^R^ vectors were produced in HEK293T packaging cells and AIF^KO^ MEFs were transduced as previously described (Perales-Clemente *et al*, 2008). Twenty-four hours after transduction, cells were selected for Geneticin (G418, Gibco) resistance.

### Growth measurements

Growth capacity in galactose-containing medium was determined by plating 5 x 10^4^ cells/well in 12-well test plates in 2 mL of the corresponding medium: high-glucose DMEM supplemented with 10% FBS, or glucose-free DMEM supplemented with 0.9 mg/mL galactose, 1 mM sodium pyruvate, and 10% FBS. Cells were incubated at 37 ^°^C for 5 days, and cell counts were recorded daily.

### Cell viability assays

Relative cell growth of the different cell lines in response to DCA compared with untreated controls was measured using Mossman’s MTT assay. Briefly, 10^4^ cells per well were seeded in a 96-well flat-bottom plates and treated with of DCA (5 mM and 25 mM) for 72 h at 37 °C. After treatment, the medium was replaced with a fresh one containing 1 mg/mL MTT, and cells were incubated for 4 h in a humidified atmosphere at 37 °C. Formazan crystals were solubilized with DMSO, and absorbance was measured at 570 nm (OD_570_) using a EZ Read 400 microplate reader (Biochrom). Results are expressed as perecentiages relative to untreated controls. All experiments were performed in at least technical and biological triplicates.

### Oxygen consumption measurements

Endogenous and maximal O_2_ consumption in intact cells were measured using an oxytherm Clark-type electrode (Hansatech) as formerly described (Hofhaus *et al*, 1996) with small modifications (Acín-Pérez *et al*, 2003).

### Protein extraction and electrophoresis

To obtain total cell protein extracts, cells were collected from 60 mm-diameter culture plates, washed twice with PBS and resuspended in RIPA buffer (50 mM Tris-HCl pH 7.4, 5 mM EDTA, 1% Triton X-100, 0.5% Sodium Deoxycholate, 50 mM NaCl) containing 1x cOmplete^TM^ Protease Inhibitor Cocktail (Roche). Steady-state levels of hAIF or CHCHD4 were estimated by SDS-PAGE electrophoresis of total cell proteins (60 μg) followed by western blotting. Samples were resolved on 12% polyacrylamide gels and electroblotted onto PVDF membranes (Amersham, Cytiva).

### Native polyacrylamide electrophoresis

Mitochondria were isolated from cultured cell lines according to Schägger (1995) (Schägger, 1995), with some modifications (Acín-Pérez *et al*, 2008). Digitonin-solubilized mitochondrial proteins (100 µg) were separated by native PAGE. Respiratory supercomplexes assembly was analysed by BN-PAGE using commercial 3-12% acrylamide native gradient gels (Novex, Thermo Fisher Scientific). To determine the ability of AIF variants to dimerize in the presence of NADH, digitonin-solubilized mitochondria were separated by CN-PAGE (4-16% acrylamide gradient) using a cathode buffer containing 0.02 % n-dodecyl β-D-maltoside and 0.05 % sodium deoxycholate (hrCNE-1) (Wittig & Schägger, 2008, 2009). When indicated, samples were preincubated with 0.4 mM NADH at room temperature for 15 min, prior to gel loading.

### Immunological techniques

Following SDS-PAGE or BN/CN-PAGE electrophoresis, proteins were electroblotted onto Amersham Hybond–P PVDF membranes (Cytiva). Primary antibodies included anti-AIF (Sigma-Aldrich), anti-CHCHD4 (Proteintech), anti-β-actin (Sigma-Aldrich), as well as specific antibodies against complex I (anti-NDUFA9; Invitrogen), complex II (anti-SDHA, 70 kDa subunit; Invitrogen), complex III (anti-UQCRC1; Invitrogen) and complex IV (anti-MT-CO1; Invitrogen). Horseradish peroxidase (HRP)-conjugated antibodies (anti-mouse or anti-rabbit; Invitrogen) enabled chemiluminiscent signal detection using the Pierce ECL Western Blotting Substrate (Thermo Fisher Scientific). Chemiluminescent signals from SDS-PAGE and BN/CN-PAGE immunoblots were captured and quantified using an Amersham Imager 600 (Cytiva) digital imaging system (ImageQuant TL, Cytiva). For immunofluorescence, cells were seeded on coverslips and incubated with 200 nM MitoTracker^TM^ Red FM (Invitrogen) at 37 °C for 30 min. Anti-AIF primary antibody (Sigma-Aldrich) and Alexa Fluor 488-conjugated goat anti-rabbit IgG secondary antibody (Invitrogen) were used for immunostaining. Immunofluorescence images were captured on a Axiovert 200M (Zeiss) fluorescence microscope using a high-resolution camera and analyzed using MetaMorph microscopy automation and image analysis software (Molecular Devices)

### Mitochondrial Superoxide production and mitochondrial mass analysis

For mitochondrial ROS production and mitochondrial mass measurement, cells were incubated with MitoSOX red (5 μM, Invitrogen) and MitoTracker^T^Green FM (200 nM, Invitrogen) for 30 min at 37 ^°^C in the dark, before assessment in a FACSCalibur (BD Biosciences) cytometer. Data were analyzed using FlowJo Software (BD Biosciences).

### Quantitative PCR determination of mRNA transcripts

Total cellular RNA was isolated from 5×10^6^ cells using TRIzol reagent (Invitrogen). One microgram of total RNA was revere-transcribed to cDNA with the Transcriptor First Strand cDNA Synthesis Kit (Roche). Transcripts for hAIF and CHCHD4 were quantified by qPCR using a LightCycler System (Roche) with the LightCycler Fast-Start DNA Master^PLUS^ SYBR Green I Kit (Roche). Amplification was carried out using gene-specific primers for h*AIF* and *Chchd4*, following the manufacturer’s recommendations. Transcript levels were normalized to *Actb* mRNA (NM_007393). Primer sequences are listed in Appendix Table S2.

### Protein expression and purification

Recombinant human CHCHD4 (UniProtKB: Q8N4Q1), soluble mitochondrial AIF_Δ77_ and its corresponding variants, E413A/R422A/R430A (herein 3M), R422Q and H454A, were produced in *Escherichia coli* using pET28a(+) expression vector encoding a removable His_6_-tag (CACCAT) followed by a PreScission Plus protease recognition site (Novo *et al*., 2023; Romero-Tamayo *et al*., 2021). The His_6_-tag is located in the C-terminal for AIF_Δ77_ variants and the N-terminal for CHCHD4. Synthetic coding sequences were cloned into the *Nco*I and *Nde*I restriction sites of the pET-28a(+) plasmid. The engineered AIF_Δ77_ variants were generated by site-directed mutagenesis (GenScript®). AIF_Δ77_ variants were expressed in C41 (DE3) cells, whereas CHCHD4 was produced in SHuffle T7 cells. A synthetic peptide containing the first 27 N-terminal aminoacids of CHCHD4 (N-CHCHD4) was purchased from NZYTech.

AIF_Δ77_ variants were produced and purified as formerly described (Romero-Tamayo *et al*., 2021). CHCHD4 expression and purification were carried out according to Fagnani *et al* (Fagnani et al., 2024; Ferrer et al., 2026). The His_6_-tag was removed by protease digestion (10 U/mg protein) for 2 h at 37 °C and cleaved proteins were separated from affinity tags by reloading the samples onto a HisTrap affinity column. AIF concentrations were estimated spectrophotometrically using previously determined flavin extinction coefficients (Ferreira *et al*., 2014; Villanueva *et al*., 2019) or, alternatively, by quantifying flavin release upon protein denaturation with 3 M guanidinium chloride in 50 mM potassium phosphate buffer, pH 7.4. The extinction coefficients, ε_451 nm_, were 13.6, 13.4, 12.2 and 12.75 mM^-1^·cm^-^1 for the WT, 3M, H454A and R422Q variants, respectively.

CHCHD4 and N-CHCHD4 concentrations were determined using their theoretical ε_280 nm_ values (13.3 and 1.49 mM^-1^·cm^-1^, respectively), calculated with the ProtParam tool (ExPASy).

### Molecular weight determination by size-exclusion chromatography (SEC)

The oligomeric state of hAIF_D77_ variants was analyzed in the absence or presence of NADH (2 mM) and/or CHCHD4. Protein mixtures, previously incubated for 15 min at room temperature when indicated, were loaded onto a Superdex 200 10/300 GL column (Cytiva) connected to an ÄKTA go protein purification system (Cytiva). Elution profiles were acquired at a flow rate of 0.4 mL/min in 50 mM phosphate buffer, 150 mM NaCl, pH 7.4. Column calibration was performed using standard proteins in the 13.7-440 kDa range (Gel filtration calibration kit, Cytiva). Chromatographic peaks were analyzed by Gaussian fitting.

### Stabilization of cross-linked protein oligomers and electrophoretic analysis

To stabilize oligomeric species, 2 µM AIF_Δ77_ variants were incubated in 50 mM potassium phosphate buffer, pH 7.4, with a 100-fold excess of BS^3^ (homobifunctional-bis[sulfosuccinimidyl]-suberate from Pierce) cross-linker for 30 minutes at room temperature, in the absence or presence of a 100-fold excess of NADH. Samples were mixed with denaturing bromophenol blue sample buffer to stop the reaction and subsequently incubated at 100 °C for 5 minutes. Cross-linked products were resolved using 12% SDS-PAGE gels.

### Spectroscopic characterization

Absorption spectra were collected using a Cary 100 Bio spectrophotometer (Agilent Technologies). Circular dichroism (CD) measurements were carried out in a thermostated Chirascan (Applied Photophysics Ltd.) at 25 °C in 50 mM potassium phosphate buffer, pH 7.4 (150 mM ionic strength), in either the absence or presence of a 100-fold excess of NADH. Near-UV/visible CD spectra were acquired using 20 µM AIF_Δ77_ variants in a 1 cm-pathlength cuvette, whereas Far-UV CD measurements were recorded using 5 µM AIF_Δ77_ variants in a 0.1 cm-pathlength cuvette.

### Thermal denaturation assays

Thermal unfolding AIF_Δ77_ variants was monitored through FAD fluorescence emission upon excitation at 450 nm. Assays were performed using 2 µM AIF_Δ77_ variants, in the absence or presence of NADH (100-fold molar excess) and/or CHCHD4/N-CHCHD4 (1:3 molar ratio relative to AIF). Fluorescence measurements were recorded in a 1 cm-pathlength cuvette from 15 °C to 90 °C at a heating rate of 1.5 °C /min.

Experimental unfolding curves were normalized and fitted to a two-state transition unfolding model (native (N) ↔ unfolded (U)) according to (Sancho, 2013):

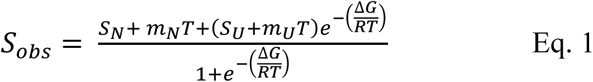

where S_obs_ corresponds to the experimental protein signal measured at a given temperature (T), and the parameters S_x_ (S_N_,S_U_) and m_x_ (m_N_, and m_U_) describe the y-intercepts at 0 K and the slopes of the pre– and post-transition baselines, respectively. The Stabilization Gibbs energy depends temperature according to 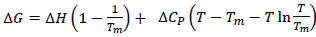, in which ΔH, T_m_ and ΔC_P_ symbolize the unfolding enthalpy, unfolding temperature and unfolding heat capacity, respectively. R represents the ideal gas constant.

### Kinetics measurements

The NADH-dependent diaphorase activity of AIF_Δ77_ variants was evaluated at 25 °C in air saturated 50 mM potassium phosphate buffer, pH 7.4, using 95 µM dichlorophenolindophenol (DCPIP, Δε_620*nm*_= 21 mM^-1^ cm^-1^) as electron acceptor (Ferreira *et al*., 2014). Assays were performed in the absence or presence of CHCHD4/N-CHCHD4 (1:3 molar ratio relative to AIF). Initial rates obtained at different NADH concentrations were fitted to the Michaelis-Menten equation to determine kinetic parameters:

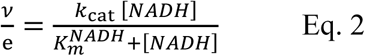

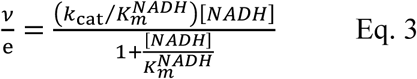

where *v/e* denotes the initial reaction rate normalized to AIF_Δ77_ enzyme concentration, *K_m_^NADH^* reflects the Michaelis constant for NADH and *k*_cat_ defines the catalytic turnover number of the enzyme under steady-state conditions. Catalytic efficiency was estimated from the *k*_cat_⁄*K_m_^NADH^* ratio.

The stability of the CTC towards molecular oxygen was assessed by fully reducing AIF_Δ77_ variants with NADH (1-1.5 molar ratio relative to AIF) in 50 mM potassium phosphate buffer, pH 7.4 according to a previous established protocol (Romero-Tamayo *et al*., 2021). Reoxidation was monitored spectrophotometrically, in the absence or presence of CHCHD4/N-CHCHD4 (1:3 molar ratio relative to AIF), using a Cary 100 Bio spectrophotometer (Agilent Technologies).

Kinetic measurements of the reductive half-reaction in AIF_Δ77_ variants were performed using an SX18.MV stopped-flow spectrophotometer (Applied Photophysics Ltd.*, Surrey, UK*). AIF_Δ77_ variants were mixed with increasing concentrations of NADH (0.03-10 mM), in the absence or presence of CHCHD4/N-CHCHD4 (1:3 molar ratio relative to AIF). Experiments were carried out in 50 mM potassium phosphate buffer, pH 7.4, at 25 °C, except for assays involving the N-CHCHD4 or the H454A_Δ77_ with CHCHD4, which were recorded at 10 °C. Data were acquired by using a photodiode array and monochromator detectors. Observed rate constants (*k*_obs_) for HT were fitted by global analysis of the spectra or by exponential fitting of single-wavelength traces according to one– or two-step reaction models using Pro-K and ProData-XD software. Averaged *k*_obs_ at each NADH concentration were subsequently non-linearly fitted to the Equation 4, which describes the formation of AIF_Δ77_ (CHCHD4):NADH complex preceding the HT event:

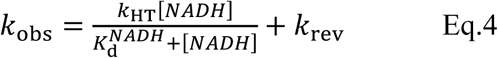

where *k*_HT_ and *k*_rev_ correspond to HT constant and its reverse reaction, respectively and *K_d_^NADH^* represents the dissociation constant of the transient AIF_Δ77_(CHCHD4):NADH complex.

### Mid-point reduction potential

Mid-point reduction potential of WT_Δ77_ and 3M_Δ77_ variants either free or in complex with the N-CHCHD4 (3-fold molar excess relative to AIF) were determined under anaerobic conditions using the xanthine/xanthine oxidase reduction system (Christgen *et al*, 2019). Briefly, reaction mixtures containing 10 µM of AIF_Δ77_ variants, 500 µM of xanthine and 2 µM of benzyl viologen (redox mediator; E_m,pH 7_ = –359 mV) were treated with multiple vacuum/argon cycles to ensure anaerobiosis. Residual oxygen was removed by addition of 10 mM of glucose and 10 U/mL of glucose oxidase after several cycles. Reduction was initiated by adding xanthine oxidase (0.002 U/mL) and the spectra were recorder spectrophotometrically every 3 min up to 3 hours. Assays were performed in 50 mM potassium phosphate buffer, pH 7.4, at 25 °C. The oxidized/reduced ratios for both dye and protein species were calculated using Equation 5 at different time points:

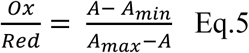

where A, A_min_ and A_max_ represent the absorbance at a given time, the dye/protein’s minimum and maximum absorbance, respectively.

The log–transformed protein and dye ratios were subsequently plotted and fitted to:

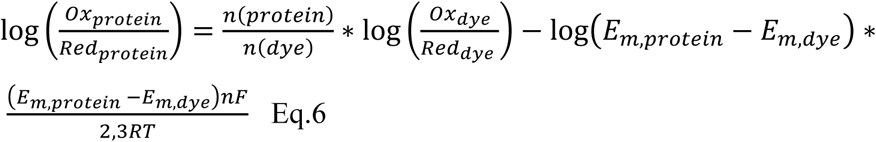

where n symbolizes the number of transferred electrons, T reflects the temperature, E_m_, defines the midpoint reduction potential and F and R correspond to the Faraday and gas constant, respectively. The E_m_ values of the AIF_Δ77_ variants and AIF_Δ77_:N-CHCHD4 complexes were finally determined using the known dye potential according to Equation 7:

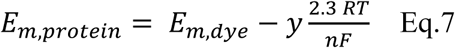

### Isothermal titration calorimetry (ITC)

Protein-protein interactions were characterized using an Auto-iTC200 (Malvern). Measurements were generally performed at 25 °C, except for WT_Δ77ox,_ which was determined at 10 °C. Typically, 100 μM CHCHD4 or N-CHCHD4 was used to titrate ∼10 μM hAIF_Δ77_ variants, in the absence or presence of a 100-fold molar excess of NADH. Prior to each experiment, samples were degassed for 3 min at 15 °C. Titrations consisted of sequential 2 μL injections of titrant solution every 150 s under constant stirring speed at 750 rpm. The binding isotherms were generated by integrating the heat effects per injection, after baseline correction, and normalizing by the amount of injected titrant. The thermodynamic parameters binding stoichiometry (N), association constant (*K*_a_) and enthalpy of binding (ΔH), were obtained through non-linear least-squares regression of the experimental data employing a single-site ligand binding model implemented in Origin 7.0 (OriginLab) software. The dissociation constant (*K*_d_), the free energy change (Δ*G*), and the entropy change (Δ*S*) were calculated from basic thermodynamic relationships.

### Data and statistical analysis

Experimental data were analyzed and shown using SigmaPlot (Systat. Software Inc.), Origin 7.0 (OriginLab), Pro-K (Applied Photophysics Ltd.) and StatView 5.0 (SAS Institute) software. Results are expressed as mean ± SD. Statistical analyses were performed using non-parametric Mann-Whitney *U* test for comparisons between two groups were performed, or the Kruskal-Wallis test for multiple comparisons. Differences were considered statistically significant at *P* ≤ 0.05. The number of technical and biological replicates is specified in the corresponding figure legends. Structural data visualization, analysis and figure generation were performed using PyMol (Delano, 2002).

## Acknowledgments

This work was funded by the Spanish State Research Agency and by FEDER (MCIN/AEI-FEDER, Grants PID2022-136369NB-I00, Grant PID2021-124354NB-I00), as well as by the Gobierno de Aragón, grant number “Grupo de Referencia E35_17R” to M.M-M., M.M, P.F.-S., R.M.-L. and P.F. and grant number “LMP220_21” to P.F.-S. and R.M.-L. The authors would like to acknowledge Servicios Generales de Apoyo a la Investigación (SAI), University of Zaragoza, for their support, as well as at BIFI-University of Zaragoza for providing instrumentation.

## Author contributions

Conceptualization, supervision and project administration: P.F and R.M-L. Data curation: P.F., R.M.-L., A.V.-C., P.F.-S and M.M. Funding acquisition: P.F., R.M.-L, M.M. and P.F.-S. Investigation and methodology: R.M.-L, S.H-H, O.S. and P. F-S (cell culture, respiration, viability analysis and western blot analysis), J.M.-B. (Flow cytometry measurements), and O.S., R.G-D., S. R-T, M.F. and P.F (protein production and molecular and biophysical characterization). Resources: S.S. (AIF^KO^ cells generation and provision). Writing original draft: P.F., R.M.-L. and O.S. Review & editing: all authors. All authors have read and approved the final version of the manuscript.

## Material availability

Material that are unique to this study are available upon request.

## Data and code availability

This paper does not report original code. Any additional information necessary to reanalyze the data reported in this paper is available from the lead contact upon request.

## Abbreviations

hAIF: human apoptosis-inducing factor
hAIF 3M: variant of hAIF harboring the E413A/R422A/R430A mutations
AIF^KO^ MEFs: a stable AIF knockout MEF cell line
hAIF^WT^: AIF^KO^ MEFs transfected with hAIF
hAIF^3M^: AIF^KO^ MEFs transfected with hAIF 3M variant
ALR: Augmenter of Liver Regeneration
CHCHD4/MIA40: Coiled-coil-helix-coiled-coil-helix domain containing 4/ Mitochondrial intermembrane space import and assembly protein 40
N-CHCHD4: peptide containing the first 27 amino acids at N-terminal of CHCHD4
CTC: Charge transfer complex
CPC: cysteine-proline-cysteine
FAD: Flavin adenine dinucleotide
HT: Hydride transfer
NADH: Reduced nicotinamide adenine dinucleotide
OXPHOS: Oxidative phosphorylation
IMS: Intermembrane space
BS^3^: Homobifunctional-bis[sulfosuccinimidyl]-Suberate
CD: Circular dichroism
ITC: Isothermal titration calorimetry
MEFs: mouse embryonic fibroblasts
GFP: Green fluorescent protein

## Legend of figures

**Figure EV1.** – Isothermal calorimetric titrations for binary interactions of AIF_Δ77_ variants with CHCHD4 and N-CHCHD4. **A-I** Calorimetric titrations of purified WT_Δ77_, 3M_Δ77_ and H454A_Δ77_ with CHCHD4 or N-CHCHD4. The upper panels show the thermograms for the interactions, whereas the lower panels show the corresponding binding isotherms with integrated heats. Data were fitted to a home-derived model for a single binding site (solid lines and closed symbols for AIF_Δ77ox_ variants and dashed lines and open symbols for AIF_Δ77red_ variants). **J** Dissection of the thermodynamic parameters for the interaction of AIF_Δ77_ variants with CHCHD4 and its N-terminal peptide in the presence or absence of NADH as assessed from experimental ITC assays. The binding Gibbs energy (Δ*G*), enthalpy (Δ*H*) and entropy (–TΔ*S*) contributions are displayed in blue, green and red bars, respectively. The reduced forms were obtained by premixing AIF_Δ77ox_ proteins with a 100-fold excess of NADH. Plots are representative of at least two independent assays.

**Figure EV2.** – CHCHD4 modulates AIF_Δ77_ redox properties. **A-C** Time course of HT from NADH to AIF_Δ77_:N-CHCHD4 complexes. Spectral evolution of flavin reduction for WT_Δ77_:N-CHCHD4 (A), 3M_Δ77_:N-CHCHD4 (B) and H454A_Δ77_:N-CHCHD4 (C) complexes. Dark red, dark cyan and blue dashed lines represent spectra of WT_Δ77ox_, 3M_Δ77ox_ and H454A_Δ77ox_, respectively, before mixing with NADH. Data were globally fitted to a single-step or a two-steps model describing the transitions from initial species A to final species B and C (see insets). Estimated *k*_obs_ is represented in each panel. Spectral evolution was recorded at 0.001, 0.004. 0.006, 0.009, 0.012, 0.017, 0.025, 0.048 and 0.25 s for WT_Δ77_:N-CHCHD4, at 0.001, 0.015, 0.028, 0.05, 0.1 and 0.3 s for 3M_Δ77_:N-CHCHD4 and at 0.001, 0.008, 0.016, 0.03 and 0.4 s for H454A_Δ77_:N-CHCHD4. **D-F** Reactivity of AIF_Δ77_ CTCs towards O_2_ for WT_Δ77_ in the presence of N-CHCHD4 (D), and for 3M_Δ77_ (E) and H454A_Δ77_ (F) in the presence of CHCHD4 and N-CHCHD4 (see insets). The AIF_Δ77rd_:NAD^+^ CTC samples were obtained by mixing AIF_Δ77ox_ proteins with a 1.5–fold excess of NADH. Dark red, dark cyan and blue lines correspond to spectra of AIF_Δ77ox_ variants in the absence (solid lines) and presence (dashed lines) of N-CHCHD4/ CHCHD4, prior to NADH addition (color code as in A-C). Black dashed dotted lines represent the WT_Δ77_ CTC half-life time.

**Table EV1.** Thermodynamic parameters for the binary interaction of AIF variants with CHCHD4.

| Sample | Titration ligand | $K_d$<br>(μM) | N | $\Delta H$<br>(kcal/mol) | $\Delta G$<br>(kcal/mol) | $-T\Delta S$<br>(kcal/mol) |
| --- | --- | --- | --- | --- | --- | --- |
| **WT$_{\Delta 77ox}$\*** | N-CHCHD4 |  |  | Binding not detected |  |  |
| **3M$_{\Delta 77ox}$** | N-CHCHD4 |  |  | Binding not detected |  |  |
| **H454A$_{\Delta 77ox}$** | N-CHCHD4 | 1.75 | 1.1 | -20.5 | -7.9 | 12.6 |
| **WT$_{\Delta 77rd}$** | N-CHCHD4 | 0.35 | 1.1 | -14.0 | -8.8 | 5.2 |
| **3M$_{\Delta 77rd}$** | N-CHCHD4 | 0.15 | 0.8 | -13.2 | -9.3 | 3.9 |
| **H454A$_{\Delta 77rd}$** | N-CHCHD4 | 0.35 | 1.0 | -15.7 | -8.9 | 6.9 |
| **WT$_{\Delta 77ox}$\*** | CHCHD4 | 7.7 | 1.2 | -26.0 | -7.0 | 19.0 |
| **3M$_{\Delta 77ox}$** | CHCHD4 |  |  | Binding not detected |  |  |
| **H454A$_{\Delta 77ox}$** | CHCHD4 | 0.9 | 0.94 | -12 | -8.3 | 3.7 |
| **WT$_{\Delta 77rd}$** | CHCHD4 | 0.8 | 1.2 | -18.1 | -8.3 | 9.8 |
| **3M$_{\Delta 77rd}$** | CHCHD4 | 0.6 | 0.7 | -12.9 | -8.5 | 4.4 |
| **H454A$_{\Delta 77rd}$** | CHCHD4 | 0.2 | 1 | -6.6 | -9.1 | -2.5 |

Values obtained from ITC assays at 25 °C in 50 mM potassium phosphate, pH 7.4. N is the estimated apparent binding stoichiometry. The thermodynamic parameters were calculated using well-known relationships: $K_d = (K_a)^{-1}$ , $\Delta G = RT \cdot \ln K_d$ and $-T\Delta S = \Delta G - \Delta H$ . Errors considered in the measured parameters ( $\pm 20\%$ in $K_d$ and $\pm 0.3$ kcal/mol in $\Delta H$ and $-T\Delta S$ ) were taken larger than the standard deviation between replicates and the numerical error after the fitting analysis. \*Values obtained from ITC assays at 10 °C in 50 mM Tris-HCl buffer, pH 8. ITC assays with 3M $_{\Delta 77ox}$ could not be performed with these conditions due to protein instability.

